# Severe Acute Respiratory Syndrome Coronavirus 2 (SARS-CoV-2) Membrane (M) Protein Inhibits Type I and III Interferon Production by Targeting RIG-I/MDA-5 Signaling

**DOI:** 10.1101/2020.07.26.222026

**Authors:** Yi Zheng, Meng-Wei Zhuang, Lulu Han, Jing Zhang, Mei-Ling Nan, Chengjiang Gao, Pei-Hui Wang

**Affiliations:** Key Laboratory of Infection and Immunity of Shandong Province, Department of Immunology, School of Basic Medical Sciences, Cheeloo College of Medicine, Shandong University, Jinan 250012, China; Advanced Medical Research Institute, Cheeloo College of Medicine, Shandong University, Jinan 250012, China; Suzhou Research Institute, Shandong University, Shandong University, Suzhou, Jiangsu 215123, China

**Author notes:** Correspondence: Chengjiang Gao or Pei-Hui Wang ( or). These authors contributed equally: Yi Zheng, Meng-Wei Zhuang.

**Keywords:** SARS-CoV-2, COVID-19, M protein, antiviral immunity, IFNs

## Abstract

The coronavirus disease 2019 (COVID-19) caused by Severe acute respiratory syndrome coronavirus 2 (SARS-CoV-2) has quickly spread worldwide and has infected more than ten million individuals. One of the typical features of COVID-19 is that both type I and III interferon (IFN)-mediated antiviral immunity are suppressed. However, the molecular mechanism by which SARS-CoV-2 evades this antiviral immunity remains elusive. Here, we report that the SARS-CoV-2 membrane (M) protein inhibits the production of type I and III IFNs induced by the cytosolic dsRNA-sensing pathway of RIG-I/MDA-5-MAVS signaling. The SARS-CoV2 M protein also dampens type I and III IFN induction stimulated by Sendai virus infection or poly (I:C) transfection. Mechanistically, the SARS-CoV-2 M protein interacts with RIG-I, MAVS, and TBK1 and prevents the formation of a multi-protein complex containing RIG-I, MAVS, TRAF3, and TBK1, thus impeding IRF3 phosphorylation, nuclear translocation, and activation. Consequently, the ectopic expression of the SARS-CoV2 M protein facilitates the replication of vesicular stomatitis virus (VSV). Taken together, the SARS-CoV-2 M protein antagonizes type I and III IFN production by targeting RIG-I/MDA-5 signaling, which subsequently attenuates antiviral immunity and enhances viral replication. This study provides insight into the interpretation of the SARS-CoV-2-induced antiviral immune suppression and sheds light on the pathogenic mechanism of COVID-19.

## Introduction

The coronavirus disease 2019 (COVID-19), caused by severe acute respiratory syndrome coronavirus 2 (SARS-CoV-2), has caused a large number of infections and fatalities worldwide, representing an acute and rapidly developing global health crisis. SARS-CoV-2 show 79.5% identity with SARS-CoV-1 and approximately 50% identity with MERS-CoV at the whole-genome level.^1–3^ SARS-CoV-2, together with SARS-CoV-1 and Middle East respiratory syndrome coronavirus (MERS-CoV), belongs to the genus betacoronavirus in the Coronaviridae family. Coronaviruses are single-stranded positive-sense RNA viruses carrying the largest genomes (26-32 kb) among all RNA virus families, with a wide range of vertebrate hosts.^4^ The coronavirus transcripts have a 5’-cap structure and a 3’ poly(A) tail. SARS-CoV-2 is an enveloped virus with a genome of ~30 kb. Upon entry into host cells, the viral genome is used as the template for replication, transcription, and the synthesis of positive-sense genomic RNA (gRNA) and subgenomic RNAs. The gRNA is packaged by the structural proteins, namely, the spike, membrane (M), and envelope proteins, to assemble progeny virions. ^5^ Similar to SARS and MERS, COVID-19 may be a life-threatening disease, which typically begins with pneumonia.^3^ SARS-CoV-1 has infected approximately 8,000 people with an ~11% fatality rate worldwide during 2002-2003, and MERS-CoV has infected ~2,500 people with a ~36% fatality rate since 2012;^6^ after its outbreak in December 2019, SARS-CoV-2 has infected 14,974,446 individuals and caused 617,254 deaths as of July 22, 2020, according to the COVID-19 Dashboard by the Center for Systems Science and Engineering (CSSE) at Johns Hopkins University (https://coronavirus.jhu.edu/map.html). One of the hallmarked clinical feathers of COVID-19 is the poor protective immunity with high levels of pro-inflammatory cytokines, suggesting that the host immune system may be involved in COVID-19 pathogenesis.^7^

Innate immunity is the first line of host defense against viruses, initiated by the recognition of pathogen-associated molecular patterns (PAMPs), such as single-stranded RNA (ssRNA), double-stranded RNA (dsRNA), and DNA, which will trigger the production of type I interferon (IFN-α/β) and type III IFN (IFN-λ 1/2/3) in infected cells. ^8,9^ The Toll-like receptor 3 (TLR3) senses dsRNA in the endosome, while the retinoic acid-inducible gene I (RIG-I) and melanoma differentiation-associated gene 5 (MDA-5) are cytosolic receptors for dsRNA.^9^ Upon recognition of dsRNA, TIR-domain-containing adapter-inducing IFN-β (TRIF) will be recruited to the cytoplasmic domain of TLR3. TRIF further associates with receptor-interacting protein 1 (RIP1), TNF receptor-associated factor 6 (TRAF6), and TANK-binding kinase 1 (TBK1). RIP1 and TRAF6 involved in the activation of the NF-κB pathway, whereas TBK1 directly phosphorylates transcription factor IRF3, which subsequently translocates to the nucleus, leading to the induction of IFNs and other pro-inflammatory cytokines.^9,10^ When viruses enter into host cells, the viral dsRNA is recognized by RIG-I/MDA-5, which will initiate an antiviral signaling cascade by interacting with mitochondrial antiviral signaling (MAVS, also called IPS-1/VISA/Cardif). MAVS then activates IκB kinase α/β (IKK) and TBK1/IKKε, which in turn activates the transcription factors NF-κB and IRF3, respectively, to induce IFNs and other pro-inflammatory cytokines. ^11^ The cytosolic DNA sensor cyclic GMP-AMP (cGAMP) synthase (cGAS) can recognize dsDNA and produce 2’-3’ cGAMP which can bind to the stimulator of interferon genes (STING) and then activate TBK1 and IRF3, leading to IFN production.^12^ The binding of the type I or III IFNs to their specific receptors, the type I IFN receptor (IFNAR) and the type III IFN receptor (IFNLR), respectively, triggers the activation of the receptor-associated Janus kinase 1 (JAK1)/tyrosine kinase 2 (TYK2), which stimulates the phosphorylation of STAT1 and STAT2.^9,13^ JAK2 also participates in type III IFN-induced STAT phosphorylation.^14^ The activated STAT1/STAT2 heterodimers associate with IRF9 to form IFN-stimulated gene factor 3 (ISGF3), which in turn translocates into the nucleus and binds the IFN-stimulated response element (ISRE) in gene promoters, thus driving the expression of IFN-stimulated genes (ISGs) that confer antiviral abilities on host cells. Type I and III IFNs induce a similar ISG signature, with type I IFN signaling leading to a more rapid induction and decline in ISG expression.^9,13^

SARS-CoV-2 is a novel emerging coronavirus that is a global threat. How this virus is recognized by the innate immune system is currently unknown. However, studies from other coronaviruses indicated that RIG-I/MDA5 participates in the sensing of coronaviruses.^15^ TLR3 has been shown to be involved in the sensing of SARS-CoV-1 in a mouse model.^8^ SARS-CoV-2 has similar replication intermediates containing dsRNAs that can serve as RIG-I/MDA5 or TLR3 ligands. Therefore, SARS-CoV-2 is likely detected by these dsRNA sensors, which would induce the production of IFNs and other pro-inflammatory cytokines.^8^

Host antiviral immunity may provide the selective pressure on viruses, which has thus resulted in distinct strategies in viruses, including coronaviruses, for counteracting IFN responses. The viral-encoded IFN antagonists are common strategies to evade host antiviral immunity.^8^ SARS-CoV-1-encoded proteins, such as nonstructural protein 1, the papain-like protease domain in NSP3, ORF3b, ORF6, M protein, and the nucleocapsid protein, have been attributed with being the cause of the antagonism of the IFNs and ISGs.^8^ The common respiratory virus, influenza A virus, also encodes the IFN antagonist nonstructural protein 1, which blocks the initial detection by the PRR through binding and masking aberrant RNA produced during infection.^6^ The generation of IFN antagonists by viruses is a common strategy to evade host antiviral immunity.^8^ One of the striking clinical features of COVID-19 is that antiviral immunity is drastically impaired.^6,16^ SARS-CoV-2 infection induces low levels of type I and III IFNs with a moderate ISG response.^6^ Type I and III IFNs have been supplied or combined together with other drugs to treat patients with COVID-19, which is shown to be effective in suppressing SARS-CoV-2 infection and may effectively prevent COVID-19.^17–20^ In Vero cells, the replication of SARS-CoV-2 is dramatically reduced when pretreated with recombinant IFN-λ or IFN-β. SARS-CoV-2 has been shown to be more sensitive than many other human pathogenic viruses, including SARS-CoV-1.^19,21^ Treatment with type I and III IFNs can dramatically inhibit SARS-CoV-2 replication in primary human airway epithelial cells with the corresponding induction of ISGs.^22^ The role of type III IFNs in the suppression of virus replication has been highlighted.^23,24^ Type III IFN shows a better treatment option than type I IFN for influenza A virus-induced disease for its induction of antiviral immunity without the pro-inflammatory responses.^25,26^ However, in some cases, the type I and III IFN treatment promote the replication of SARS-CoV-2, and this phenotype was more pronounced in certain SARS-CoV-2 isolates.^27^ Two recent studies showed that ACE2, the cellular receptor of SARS-CoV-2, may act as a novel ISG upregulated by type I IFNs to facilitate SARS-CoV-2 infection. ACE2 is indeed upregulated by IFN treatment and SARS-CoV-2 infection, which has been observed in COVID19 patients.^28,29^ Therefore, the interactions between SARS-CoV-2 and the host type I and III IFN responses merit extensive investigations.

Overall, in most conditions, the host type I and III IFNs play an important role in restricting the infection and replication of SARS-CoV-2. To combat this, SARS-CoV-2 likely encodes multiple viral proteins to antagonize IFN responses akin to other coronaviruses. Because of its recent emergence, there is a paucity of information regarding the interaction between host antiviral immunity and SARS-CoV-2 infection. Thus, dissection of the evasion mechanism of type I and III IFN responses by SARS-CoV2 will contribute to the understanding of the pathogenesis of COVID-19 and a corresponding treatment. Here, we report that the SARS-CoV-2 M protein acts as an antagonist of both types I and III IFNs by affecting the multi-protein complex formation of RIG-I-MAVS-TRAF3-TBK1 signalosome. The SARS-CoV-2 M protein inhibits type I and III IFN production by Sendai virus (SeV) infection, poly (I:C) transfection, and the overexpression of the RIG-I/MDA-5 pathway signaling molecules. The ectopic expression of the SARS-CoV-2 M protein facilitates the replication of vesicular stomatitis virus (VSV). This study reveals a previously undiscovered mechanism of SARS-CoV-2 in evading host antiviral immunity, which may partially explain the clinical features of impaired antiviral immunity in COVID-19 patients and provide insights into the viral pathogenicity and treatment.

## Materials and methods

### Cell culture and transfection

HEK293, HEK293T, HeLa, and Vero-E6 cells were cultured in Dulbecco’s modified Eagle’s medium (DMEM, Gibco, USA) with 10% heat-inactivated fetal bovine serum (FBS, Gibco, USA). All cells were cultured at 37°C in a humidified incubator with 5% CO_2_. The plasmids were transfected into HEK293, HEK293T, and HeLa cells by Polyethylenimine ‘Max’ (Polysciences, Inc., Germany). Poly (I:C) (Sigma P1530, USA) was transfected into cells using Lipofectamine 2000 (Thermo Fisher, USA) as described previously. ^29^

### Antibodies and reagents

Rabbit anti-DYKDDDDK Tag (D6W5B), rabbit anti-IRF3 (D83B9), rabbit anti-pIRF3 (4D46), rabbit anti-TBK1 (3031S), rabbit anti-pTBK1 (D52C2), and rabbit anti-TRAF3 were from Cell Signaling Technology (USA); Mouse anti-MAVS was from Santa Cruz Biotechnology (USA); Mouse anti-actin and rabbit anti-calnexin were from proteintech (Wuhan, China); Mouse anti-Flag M2 was from Sigma Aldrich (USA); Mouse anti-Myc (9E10) Ab was from Origene (USA); Rabbit anti-GM130 was from Abcam (United Kingdom); Mouse anti-HA was from MDL biotech (China). Protein A/G beads were from Santa Cruz Biotechnology, and the anti-Flag magnetic beads were from Bimake (USA). Alexa Fluor 488 goat anti-rabbit IgG secondary antibody, Alexa Fluor 568 goat anti-mouse IgG secondary antibody, Alexa Fluor 488 goat anti-mouse IgG secondary antibody, and Alexa Fluor 568 goat anti-rabbit IgG secondary antibody were from Thermo Fisher Scientific (USA).

### Constructs and plasmids

The RIG-I, RIG-IN, MDA-5, MAVS, TBK1, IKKε, IRF3-5D, TRIF, and STING genes were cloned into pcDNA6B-Flag, pcDNA6B-Myc, pcDNA6B-V5, pCAG-Flag or pCMV-HA-N expression vectors using standard molecular cloning methods as described in our previous publications.^30–32^ The IFN-β luciferase reporter plasmid pGL3-IFN-β-Luc vector was constructed in our previous study.^33,34^ The IFNλ1 luciferase reporter plasmid pGL3-IFNλ1-Luc was constructed by inserting the 1000-bp promoter region of human IFNλ1 (nucleotides −1000 to +1, with the translation start site set as 1) into pGL3-Basic (Promega, USA) according to methods outlined in previous studies.^13,34^ The ISG luciferase reporter plasmid pISRE-Luc vector was purchased from Clontech (USA). The SARS-CoV-2 M protein gene (NCBI access No. MN908947) was synthesized (GENERAL BIOL, China) and subcloned into the pCAG-Flag expression vector. The primers used in plasmid construction are listed in Supplemental Table 1.

### Real-time quantitative PCR

Total RNA isolated with TRIzol reagent (Invitrogen) was reverse-transcribed into first-strand cDNA with the HiScript III 1st Strand cDNA Synthesis Kit with gDNA wiper (Vazyme, China) as per the manufacturer’s instructions. Real-time quantitative PCR (RT-qPCR) assays were performed using the SYBR Green-based RT-qPCR kit UltraSYBR Mixture (CWBIO, China) with a Roche LightCycler96 system as per the manufacturer’s instructions. The relative abundance of the indicated mRNA was normalized to that of GAPDH. A comparative C_T_ method (ΔΔC_T_ method) was used for the calculation of fold change in gene expression as described previously.^32,35^ The primers used in RT-qPCR analysis are listed in Supplemental Table 1.

### Luciferase reporter assays

To determine the activation of the luciferase reporters, including IFN-β-Luc, IFNλ1-Luc, and ISRE-Luc, by the proteins indicated in each experiment, a dual-luciferase reporter assay was performed as described in our previous studies.^32,35^ Briefly, approximately 0.5 x 10^5^ HEK293T cells were seeded in 48-well plates and transfected 12 hours later with the luciferase reporter plasmid and the expression vector plasmids of RIG-I, RIG-IN, MDA-5, MAVS, TBK1, IKKε, IRF3-5D, TRIF, and STING, alone or together with the plasmid expressing the SARS-CoV-2 M protein, as indicated in the experiments. The pRL-TK Renilla luciferase reporter (Promega, USA) was cotransfected to normalize the transfection efficiency and serve as an internal control. Thirty-six hours after transfection, the cells were harvested and lysed to assess the luciferase activities using the Dual-Luciferase Reporter Assay Kit (Vazyme, China) according to the manufacturer’s protocol. The luciferase activity was measured in a Centro XS3 LB 960 microplate luminometer (BERTHOLD TECHNOLOGIES, Germany). The relative luciferase activity was calculated by normalizing firefly luciferase activity to that of Renilla luciferase.

### Viruses and infection

VSV-enhanced green fluorescent protein (eGFP) and SeV were used to infect HeLa, HEK293, or HEK293T cells as described in our previous publications.^30–32^ Briefly, before infection, prewarmed serum-free DMEM medium at 37°C was used to wash the target cells, after which the virus was diluted to the desired multiplicity of infection (MOI) in serum-free DMEM and incubated with the target cells for 1-2 hours. At the end of the infection, the virus-medium complexes were discarded, and DMEM containing 10% FBS was added.

### Coimmunoprecipitation and immunoblotting

For coimmunoprecipitation assays, HEK293T cells were collected 24 hours after transfection and lysed in lysis buffer [1.0% (v/v) NP-40, 50 mM Tris-HCl, pH 7.4, 50 mM EDTA, 0.15 M NaCl] supplemented with a protease inhibitor cocktail (Sigma, USA) and a phosphatase inhibitor cocktail (Sigma, USA) as described in our previous publications.^30,31^ After centrifugation for 10 min at 14,000 g, the supernatants were collected and incubated with the indicated antibodies, followed by the addition of protein A/G beads (Santa Cruz, USA), anti-Flag magnetic beads (Bimake, USA), or anti-Myc magnetic beads (Bimake, USA). After incubation overnight at 4°C, the beads were washed four times with lysis buffer. The immunoprecipitates were eluted by boiling with 2×SDS loading buffer containing 100 mM Tris-HCl pH 6.8, 4% (w/v) SDS, 20% (v/v) glycerol, 0.2% (w/v) bromophenol blue, and 1% (v/v) 2-mercaptoethanol.

For immunoblot analysis, the M-PER Protein Extraction Reagent (Pierce, USA) supplemented with a protease inhibitor cocktail (Sigma, USA) was used to lyse the cells. The protein concentrations in the extracts were measured with a bicinchoninic acid assay (Pierce, USA) and were made equal in the different samples with extraction reagent. Total cell lysates or immunoprecipitates prepared as described above were electrophoretically separated by SDS-PAGE, transferred onto a polyvinylidene difluoride membrane (Millipore, Germany), blocked with 3% (w/v) bovine serum albumin (BSA), probed with the indicated primary antibodies and corresponding secondary antibodies, and visualized with the ECL Western Blotting Detection Reagents (Pierce, USA).

### Confocal immunofluorescence microscopy

Confocal immunofluorescence microscopy studies were performed as described in our previous publications.^30,31^ Briefly, HeLa cells were grown on 12-well slides one day before transfection with the indicated plasmids. Transfected or infected HeLa cells were then fixed in 4% paraformaldehyde, permeabilized with 0.2% Triton X-100, and blocked with phosphate-buffered saline (PBS) containing 5% horse serum and 1% BSA. The fixation, permeabilization, and blocking buffers were all purchased from Beyotime Biotechnology (China). The cells were then reacted with the indicated primary antibodies at 4°C overnight, rinsed, and reacted with corresponding secondary antibodies (Invitrogen, USA). The nuclei were counterstained with DAPI (Abcam, USA). Images were taken with a Zeiss LSM780 confocal microscope (Germany).

### Viral plaque assays

Viral plaque assays were performed on Vero-E6 cells to measure the titer of VSV-eGFP as described in our previous study.^32^ Briefly, Vero cells were seeded on 24-well plates. The next day, the cells at approximately 100% confluency were infected with serial dilutions of VSV-eGFP for 30 min. After infection, the medium was replaced with DMEM containing 0.5% agar and 2% FBS. After the agar overlay turned solid, the cells were cultured for 20-24 hours, after which the cells were fixed with a 1:1 methanol-ethanol mixture for 30 min. After removing the solid agarose-medium mix, the cells were stained with 0.05% crystal violet, and the plaques on the monolayer were then counted to calculate the virus titer.

### Bioinformatics analysis

The transmembrane motifs were predicted with the TMHMM server version 2.0 (http://www.cbs.dtu.dk/services/TMHMM/).

### Statistics

The results are representative of three independent experiments and are presented as the mean ± SEM. For statistical analysis, two-tailed unpaired Student’s t-tests were performed by GraphPad Prism 8.0 and Microsoft Excel. The P values are presented within each figure or figure legend. In all cases, a value of P□<□0.05 was considered to be statistically significant.

## Results

### The SARS-CoV-2 M protein inhibits type I and III IFN induction by SeV and poly (I:C)

To explore whether the SARS-CoV-2 M protein affects type I and III IFN production, HEK293T cells expressing the SARS-CoV-2 M protein were infected with SeV or transfected with a dsRNA mimic, poly (I:C). The expression of IFN-β, IFNλ1, and two ISG, ISG56 and CXCL10, were determined by RT-qPCR. The results indicated that both SeV infection and poly (I:C) transfection strongly stimulated the expression of IFN-β, IFNλ1, ISG56, and CXCL10 in the control HEK293T cells (Fig. 1). In HEK293T cells expressing the SARS-CoV-2 M protein, the induction of IFN-β, IFNλ1, ISG56, and CXCL10 by SeV and poly (I:C) was significantly suppressed compared with that in the HEK293T cells transfected with empty vector (Fig. 1). Therefore, the SARS-CoV-2 M protein inhibits the SeV- and poly (I:C)-induced upregulation of IFN-β, IFNλ1, ISG56, and CXCL10, suggesting that it is involved in antagonizing the IFN response.

**Figure 1.**
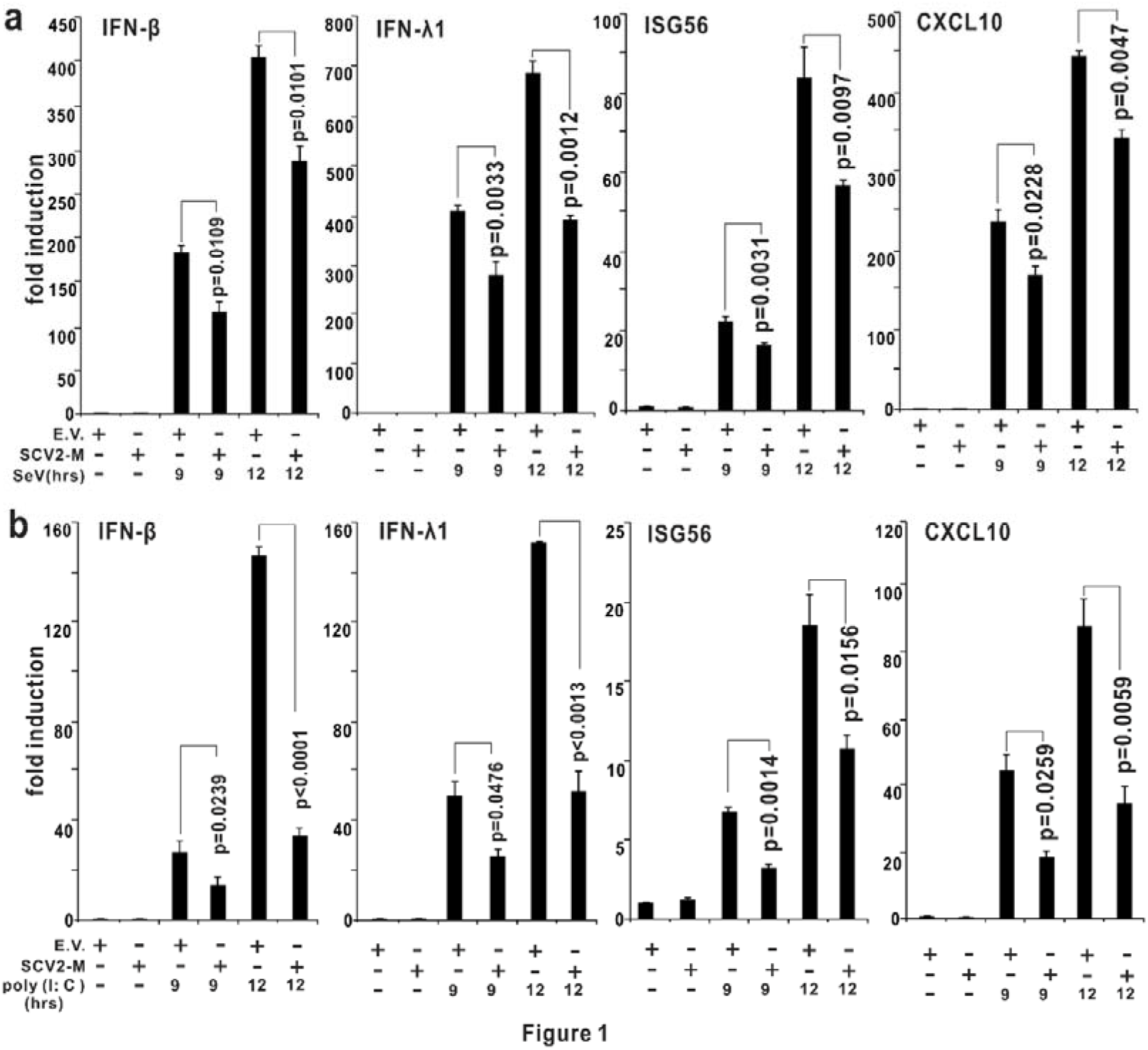
The SARS-CoV-2 M protein inhibits the induction of IFN-β, IFNλ1, ISG56, and CLXL10 by SeV infection and poly (I:C) transfection. HEK293T cells cultured in 24-well plates (0.8-1 x 10^5^ per well) were transfected with a pcDNA6B empty vector (E.V., 500 ng) or an SARS-CoV-2 M protein-expressing plasmid (500 ng). Twenty-four hours after transfection, the cells were stimulated with SeV infection (50 HA/mL) or poly (I:C) (1000 ng/mL) transfection as indicated, and at 9 and 12 hours after stimulation, the cells were harvested for RNA extraction and subsequent RT-qPCR analysis. Three independent biological replicates were analyzed; the results of one representative experiment are shown, and the error bars indicate SEM. The statistical significance is shown as indicated. SARS-CoV-2 M protein, SCV2-M; hours, hrs.

### The SARS-CoV-2 M protein dampens the cytosolic dsRNA-sensing pathway mediated by RIG-I/MDA-5 signaling

RIG-I and MDA-5 are cytosolic dsRNA sensors, which participate in the recognition of SeV- and poly (I:C) and the subsequent induction of type I and III IFNs. To further confirm the inhibitory effect of the SARS-CoV-2 M protein on type I and III IFN expression, we employed luciferase reporters of type I and III IFNs and ISGs to determine whether the SARS-CoV-2 M protein also interferes with the activation of the cytosolic RNA-sensing pathway induced by the overexpression of the RIG-I/MDA-5 pathway components. The results from IFN-β luciferase reporter (IFN-β-Luc) assays showed that overexpression of the SARS-CoV-2 M protein significantly suppressed the activities of IFN-β-Luc induced by RIG-IN (an active form of RIG-I), MDA-5, MAVS, TBK1, and IKKε but had no effect on IRF3-5D (an active form of IRF3)-, TRIF-, or STING-induced IFN-β-Luc activation (Fig. 2). Similarly, we observed that the SARS-CoV-2 M protein decreased the activities of IFNλ1-Luc (IFNλ1 luciferase reporter) and ISRE-Luc (ISG luciferase reporter) induced by RIG-IN, MDA-5, MAVS, TBK1, and IKKε instead of IRF3-5D, TRIF, or STING (Fig. 2). Consistently, in HEK293T cells expressing the SARS-CoV-2 M protein, the induction of ISG56 by RIG-I, MDA-5, MAVS, TBK1, and IKKε rather than IRF3-5D, TRIF, or STING was significantly restrained (Supplemental Fig. 1). Taken together, the SARS-CoV-2 M protein can impair the activation of the RIG-I/MDA-5-dependent cytosolic dsRNA-sensing pathway but does not affect the TLR3-TRIF signaling-mediated endosome dsRNA-sensing pathway or the cGAS-STING-mediated cytosolic DNA-sensing pathway. Because TBK1 is also downstream of TLR3-TRIF and cGAS-STING signaling, the SARS-CoV-2 M protein might thus inhibit the dsRNA-induced IFN production at the step upstream of TBK1.

**Figure 2.**
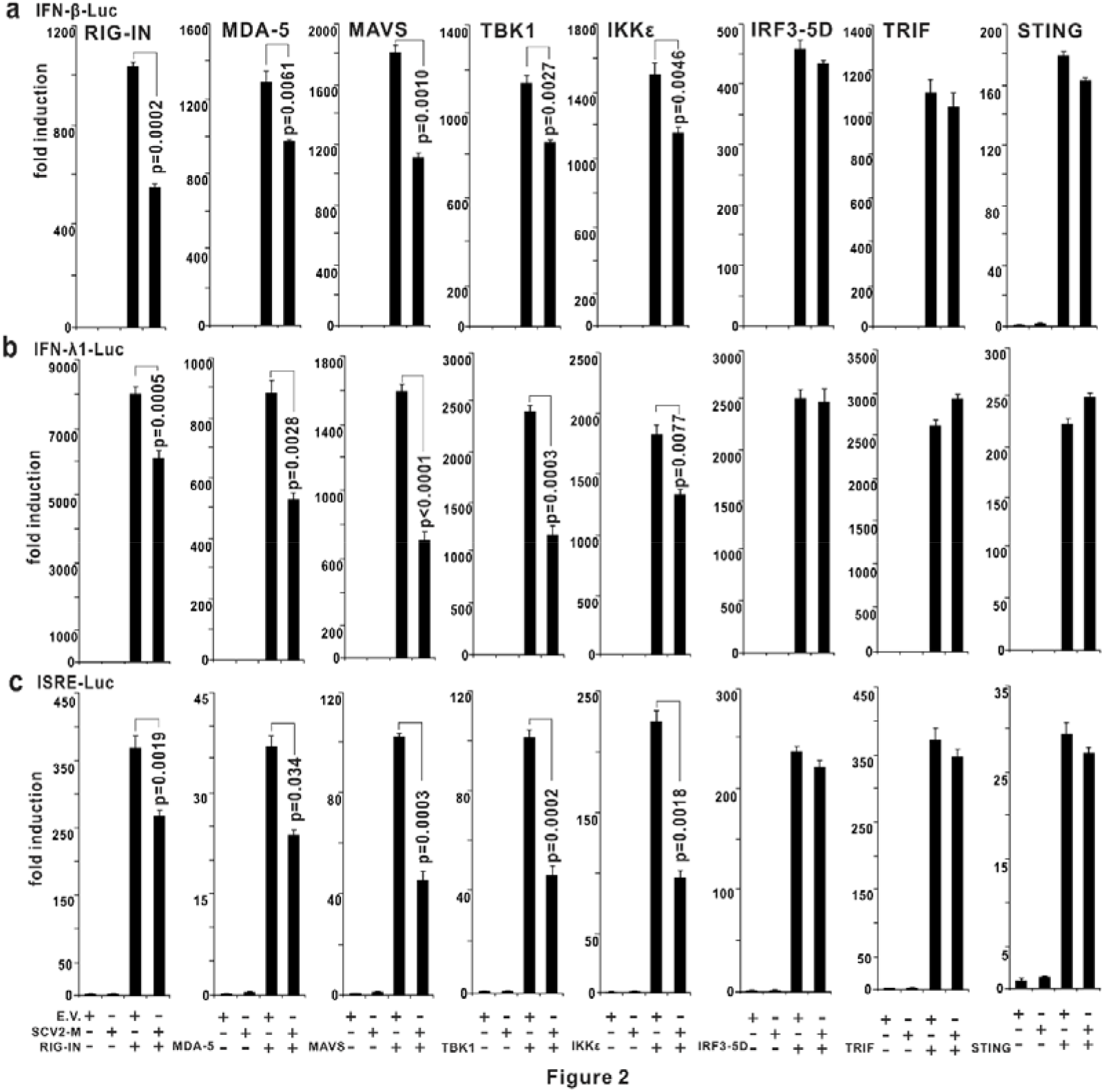
The SARS-CoV-2 M protein suppresses the luciferase reporters of type I and III IFNs and ISGs. The pcDNA6B empty vector (E.V.) and the SARS-CoV-2 M protein-expressing plasmid (100 ng) were transfected with the indicated combinations of plasmids expressing RIG-IN (100 ng), MDA-5 (100 ng), TBK1 (100 ng), IKK: (100 ng), IRF3-5D (100 ng, an active form of IRF3), TRIF (100 ng, component of TLR3-TRIF pathway), or STING (100 ng, component of cGAS-STING pathway) into HEK293T cells cultured in 48-well plates (0.5 x 10^5^ per well). Plasmids containing IFN-β-Luc (45 ng, the IFN-β luciferase reporter), IFNλ1-Luc (45 ng, the IFNλ1 luciferase reporter), or ISRE-Luc (45 ng, the IFN-stimulated response element luciferase reporter) were also transfected for indicating the activation of type I IFNs, type III IFNs, or ISGs, respectively. The pRL-TK (5 ng) was transfected into each well as an internal control. The pcDNA6B empty vector was used to balance the total amount of plasmid DNA in the transfection. Dual-luciferase assays were performed 36 hours after transfection. Three independent biological replicates were analyzed; the results of one representative experiment are shown, and the error bars indicate SEM. The statistical significance is shown as indicated. SARS-CoV-2 M protein, SCV2-M.

### Subcellular localization of the SARS-CoV-2 M protein

The SARS-CoV-2 M protein is predicted to possess three transmembrane motifs at the N terminus (Supplementary Fig. 2). Therefore, it is of interest to determine the subcellular localization of the SARS-CoV-2 M protein. To address this issue, a Flag-tagged SARS-CoV-2 M protein was overexpressed in HeLa cells, reacted with the Flag antibody and then stained with a fluorescence-labeled secondary antibody. The subcellular localization of the SARS-CoV-2 M protein was observed by confocal microscopy. The results indicated that the SARS-CoV-2 M protein showed almost no localization with mitochondria markers but was primarily localized to the endoplasmic reticulum (ER) and Golgi (Fig. 3a-c). The mitochondria, ER, and Golgi are important platforms for the multi-protein complex formation and signaling transduction of the RIG-I/MDA-5 pathway, and therefore we assessed the colocalization of the SARS-CoV-2 M protein with RIG-I, MDA-5, MAVS, TRAF3, and TBK1. The results showed that the SARS-CoV-2 M protein showed a strong colocalization signal with TBK1 but had a partial colocalization signal with RIG-I, MDA-5, and MAVS (Fig. 3d-h).

**Figure 3.**
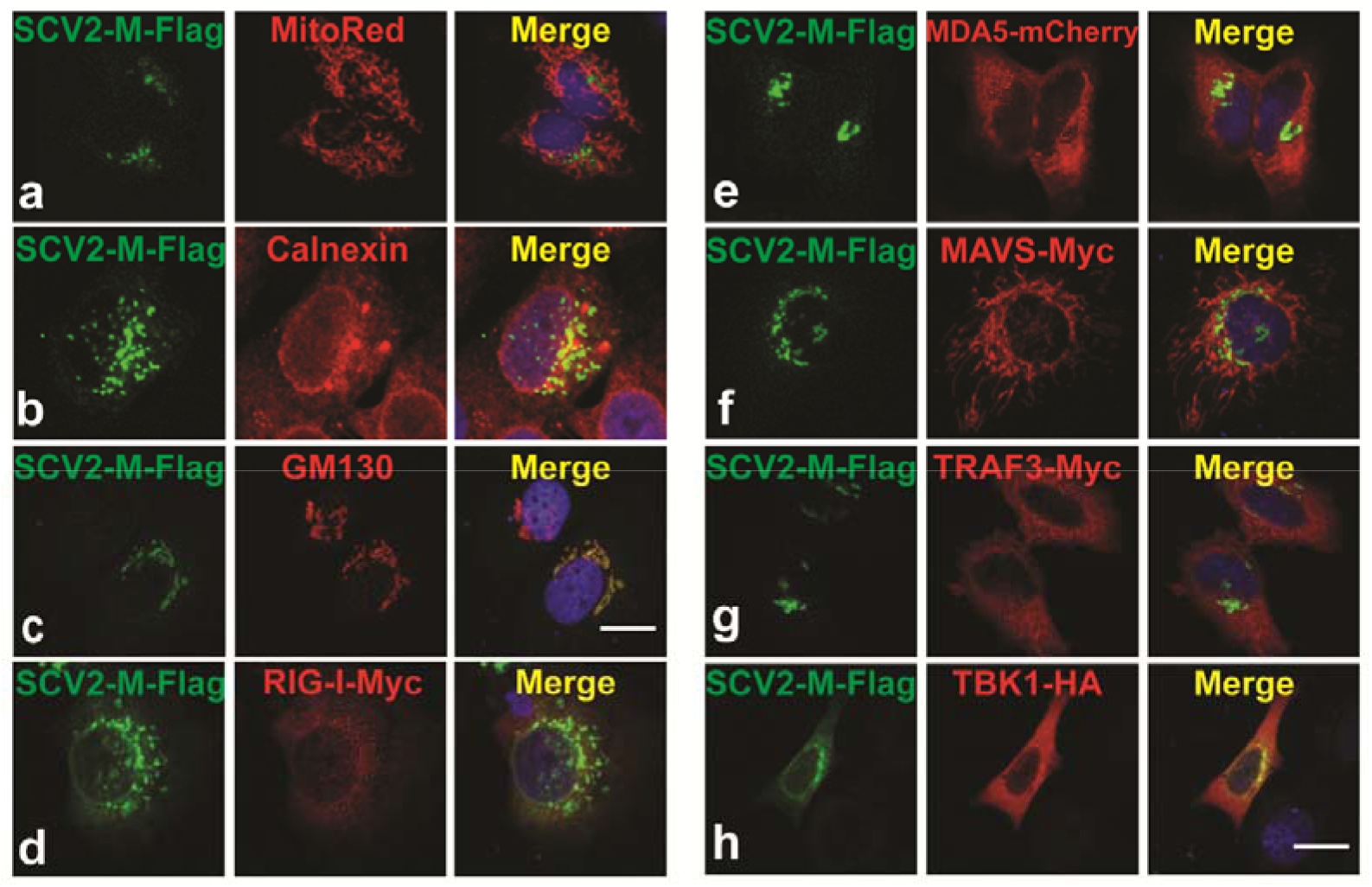
Subcellular localization of the SARS-CoV-2 M protein. HeLa cells seeded on 12-well coverslips were transfected with the indicated plasmids. Twenty hours later, the cells are fixed, blocked, and then incubated with a rabbit anti-Flag antibody and a mouse antibody against the corresponding organelle marker (**a-c**) or the indicated protein (**d, f-h**). Subsequently, the proteins were stained with a fluorescence-labeled secondary antibody. Nucleus were visualized with DAPI (blue). Confocal imaging results are representative of two independent experiments. Scale bar, 10 μm. MitoRed, mitochondria marker; Calnexin, ER marker; CM130, Golgi marker; SARS-CoV-2 M protein, SCV2-M.

### The SARS-CoV-2 M protein interacts with RIG-I, MDA-5, MAVS, and TBK1

To determine how the SARS-CoV-2 M protein affects IFN signaling activation, coimmunoprecipitation experiments between the SARS-CoV-2 M protein and the RIG-I-like receptor signaling molecules were performed. We observed the colocalization of the SARS-CoV-2 M protein with RIG-I, MDA-5, MAVS, and TBK1, and thus we examined the interactions between the SARS-CoV-2 M protein with RIG-I, MDA-5, MAVS, and TBK1. In HEK293T cells, the SARS-CoV-2 M protein was coexpressed with RIG-I, MDA-5, MAVS, TBK1, and IRF3 (Fig. 4), after which coimmunoprecipitation experiments were performed. We found that RIG-I, MDA-5, MAVS, and TBK1 but not IRF3 could be detected in the SARS-CoV-2 M protein immunoprecipitates (Fig. 4), and thus the SARS-CoV-2 M protein can associate with RIG-I, MDA-5, MAVS, and TBK1 rather than IRF3. Consistent with the coimmunoprecipitation results, we observed a strong colocalization between the SARS-CoV-2 M protein and TBK1 and a partial colocalization between the SARS-CoV-2 M protein and RIG-I, MDA-5, or MAVS (Fig. 3).

**Figure 4.**
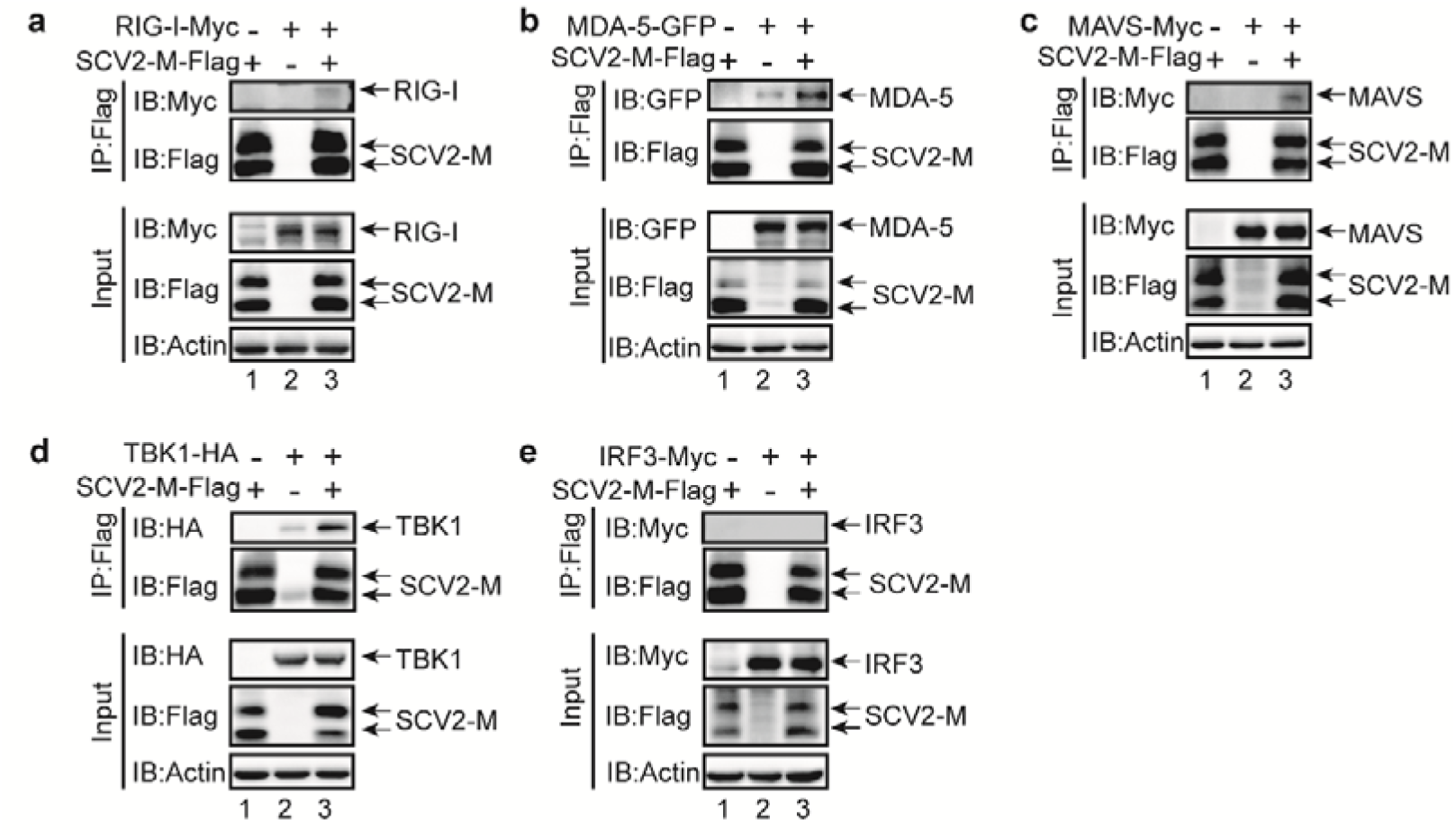
The SARS-CoV-2 M protein interacts with RIG-I (a), MDA-5 (b), MAVS (c), and TBK1 (d) but not with IRF3 (e). The HEK293T cells were transfected with the indicated plasmids for twenty-four hours before coimmunoprecipitation by the anti-Flag magnetic beads. The pcDNA6B empty vector was used to balance the total amount of plasmid DNA in the transfection. The input and immunoprecipitates were immunoblotted with the indicated antibodies. Immunoblotting results are representative of two independent experiments. SARS-CoV-2 M protein, SCV2-M.

### The SARS-CoV-2 M protein prevents RIG-I-MAVS, MAVS-TBK1, and TRAF3-TBK1 interactions and inhibits IRF3 phosphorylation

We observed the interaction between the SARS-CoV-2 M protein and RIG-I, MDA-5, MAVS, and TBK1 (Fig. 3a-d). However, an association between the SARS-CoV-2 M protein and IRF3 was unable to be detected (Fig. 3e). IRF3 activation and subsequent IFN activation relies on the assembly of a multi-protein complex containing the dsRNA sensor RIG-I/MDA-5, MAVS, TRAF3, and TBK1. Since the SARS-CoV-2 M protein can interact with RIG-I, MDA-5, MAVS, and TBK1, it of interest to investigate whether the SARS-CoV-2 M protein can affect RIG-I/MDA-5-MAVS-TRAF3-TBK1 complex formation, which is essential for IRF3 activation and IFN induction. The expression plasmids of the SARS-CoV-2 M protein and plasmids expressing RIG-I or MDA-5 were cotransfected into HEK293T cell, 24 hours later, MAVS antibodies were used to perform coimmunoprecipitation. When the SARS-CoV-2 M protein was overexpressed, the binding between RIG-I and MAVS was reduced (Fig. 5a, lanes 2 compared to lane 3); however, in the same condition, the interaction between MDA-5 and MAVS was not affected (Fig 5b, lanes 2 compared to lane 3), indicating that the SARS-CoV-2 M protein impedes the complex formation of RIG-I and MAVS but has no effect on the interaction between MDA-5 and MAVS. When using the TBK1 antibody to perform endogenous coimmunoprecipitation, MAVS was detected in the TBK1 immunoprecipitates (Fig. 5c). Moreover, when the SARS-CoV-2 M protein was overexpressed in HEK293T cells, TBK1-associated MAVS was apparently reduced compared with the cells without SARS-CoV-2 M protein expression (Fig. 5c, lanes 1 and 3 compared to lane 2; Fig. 5d, lane 4 compared to lane 2), suggesting that the SARS-CoV-2 M protein might preferentially reduce the complex formation of MAVS with TBK1. Notably, when the protein complex of TBK1 and MAVS was reduced, IRF3 phosphorylation induced by RIG-IN was correspondingly impaired (Fig 5c, lane 3 compared to lane 2). Similarly, the endogenous coimmunoprecipitation of TBK1 indicated that the overexpression of the SARS-CoV-2 M protein impeded the endogenous association between TRAF3 and TBK1 (Fig. 5d, lane 4 compared to lane 2) and thus correspondingly suppressed IRF3 phosphorylation (Fig. 5d, lane 4 compared to lane 2). Previous studies have shown that the SARS-CoV-1 M protein and MERS-CoV M protein can diminish the association between TBK1 and TRAF3 to interfere with IFN production.^36,37^ Here, we found that the overexpression of the SARS-CoV-2 M protein not only decreased the interaction between TBK1 and TRAF3 but also reduced the binding between RIG-I and MAVS and the binding between MAVS and TBK1, suggesting that the SARS-CoV-2 M protein affects the multi-protein complex formation of RIG-I-MAVS-TRAF3-TBK1 and the subsequent IRF3 phosphorylation.

**Figure 5.**
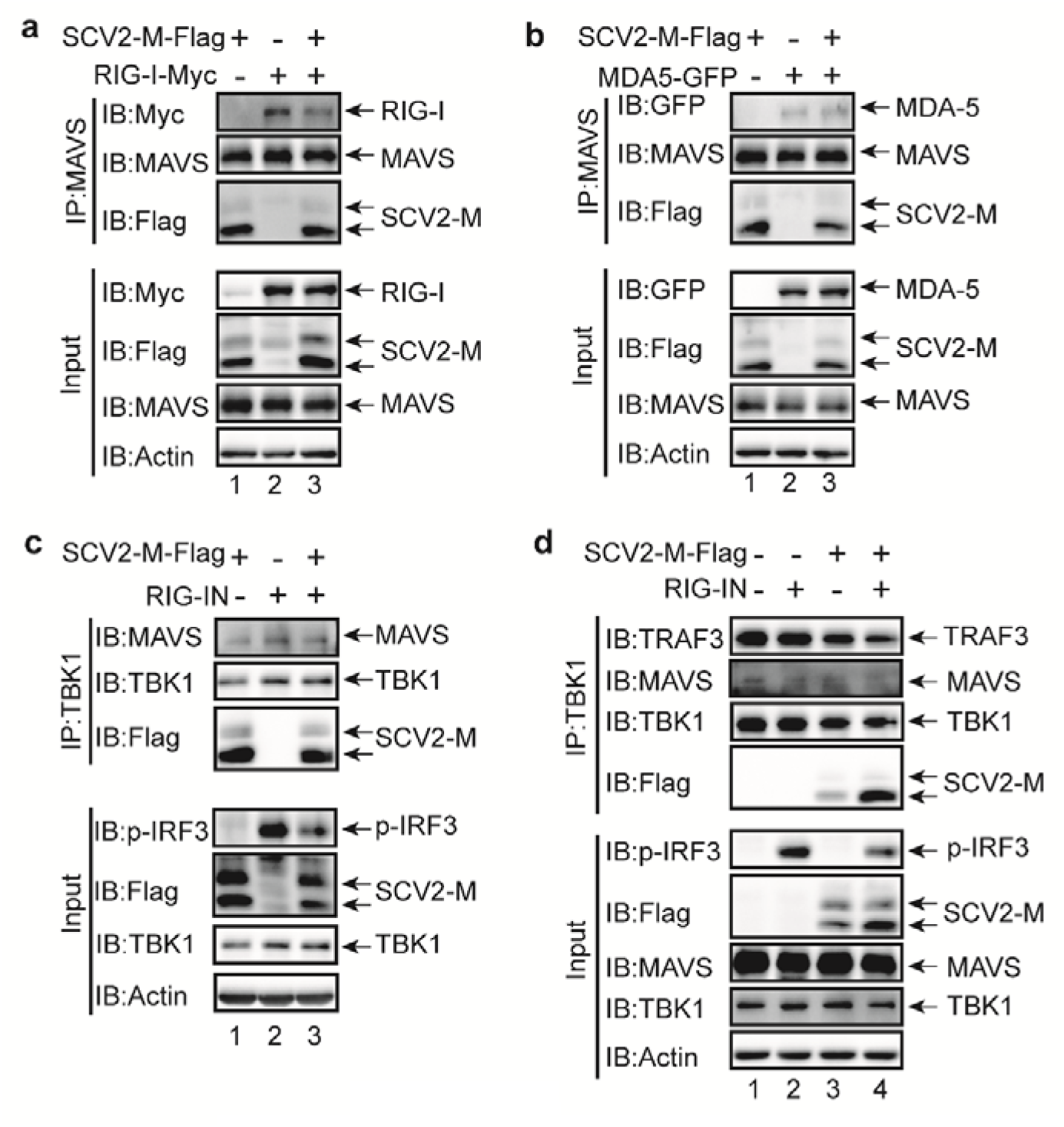
The SARS-CoV-2 M protein prevents the multi-protein complex formation involving RIG-I, MAVS, TRAF3, and TBK1. The SARS-CoV-2 M protein inhibits the RIG-I-MAVS (**a**), MAVS-TBK1 (**c**), and TRAF3-TBK1 (**d**) interactions. The HEK293T cells were transfected with the indicated plasmids for 24 h before coimmunoprecipitation by the MAVS (**a-b**) or TBK1 (**c-d**) antibody. The pcDNA6B empty vector was used to balance the total amount of plasmid DNA in the transfection. The input and immunoprecipitates were immunoblotted with the indicated antibodies. Immunoblotting results are representative of two independent experiments. SARS-CoV-2 M protein, SCV2-M.

### The SARS-CoV-2 M protein suppresses SeV-induced IRF3 phosphorylation and nuclear translocation

The phosphorylation and nuclear translocation of IRF3 is the hallmark of its activation, which is essential for type I and III IFN induction during virus infection. IRF3 phosphorylation is the prerequisite for its nuclear translocation and IFN transcription. Therefore, we next investigated the effect of the SARS-CoV-2 M protein on IRF3 phosphorylation. The results from RT-qPCR analysis (Fig. 1) and a luciferase reporter assay (Fig. 2) have indicated that the SARS-CoV-2 M protein could inhibit the induction of type I and III IFN and that the overexpression of the SARS-CoV-2 M protein reduced the IRF3 phosphorylation induced by RIG-IN (Fig. 5c, d). However, whether the SARS-CoV-2 M protein affects IRF3 phosphorylation in a real virus infection is unknown, which may contribute to our understanding of the role of the SARS-CoV-2 M protein in SARS-CoV-2 infection. Because of the lack of a biosafety level 3 laboratory, we used another RNA virus, SeV, as a surrogate for SARS-CoV-2 to perform the virus infection studies. To address the effect of the SARS-CoV-2 M protein on virus-induced IRF3 phosphorylation, control HeLa cells and HeLa cells expressing the SARS-CoV-2 M protein were infected with SeV. The immunoblotting results indicated that SeV infection could induce the phosphorylation of IRF3 in HeLa cells, while the phosphorylation of IRF3 was obviously decreased in HeLa cells expressing the SARS-CoV-2 M protein (Fig. 6a). Therefore, the SARS-CoV-2 M protein can inhibit IRF3 phosphorylation induced by SeV infection. In contrast, TBK1 phosphorylation was not affected in HeLa cells expressing the SARS-CoV-2 M protein (Fig. 6a).

**Figure 6.**
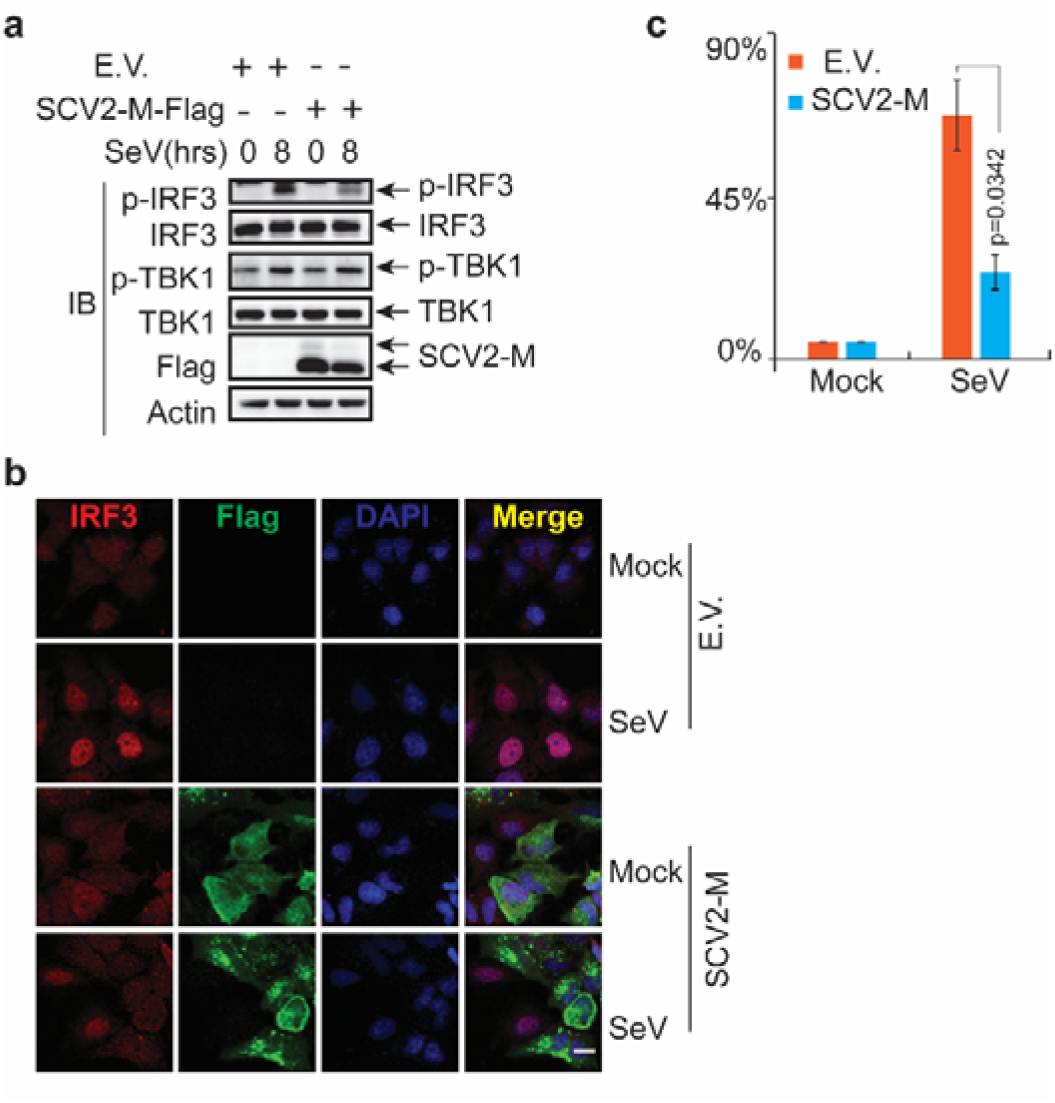
The SARS-CoV-2 M protein suppresses IRF3 phosphorylation and nuclear translocation. **a** The SARS-CoV-2 M protein affects the phosphorylation of IRF3 upon SeV infection. HeLa cells seeded on 6-well plates were transfected with the plasmid of pCAG-Flag empty vector or the plasmid of Flag-tagged SARS-CoV-2 M protein for 20 hours before infection with SeV (50 HA/mL). At the indicated time points, the cells were harvested and processed for immunoblotting with the indicated antibodies. **b** The SARS-CoV-2 M protein prevents the nuclear translocation of IRF3. HeLa cells seeded on 12-well coverslips were transfected with the plasmid of pCAG-Flag empty vector or the plasmid of Flag-tagged SARS-CoV-2 M protein for 20 hours before infection with SeV. After infection for 8 hours, the slides were harvested and processed for immunofluorescence with a mouse anti-Flag antibody and a rabbit anti-IRF3 antibody. Scale bar, 10 μm. **c** Quantification of the percentage of IRF3 in the nucleus upon SeV infection. IRF3 present in the nucleus from 50 cells within each group from (**b**) was counted and calculated. Immunoblotting and confocal imaging results are representative of two independent experiments. Empty vector, E.V.; SARS-CoV-2 M protein, SCV2-M; hours, hrs.

The phosphorylation of IRF3 is a pivotal step for its nuclear translocation to initiate the transcription of type I and III IFNs. Because we observed that the SARS-CoV-2 M protein suppressed SeV-induced IRF3 phosphorylation (Fig. 6a), we next examined the effect of the SARS-CoV-2 M protein on SeV-induced IRF3 nuclear translocation. In HeLa cells, IRF3 was evenly distributed in both the nucleus and the cytosol in the absence or presence of the SARS-CoV-2 M protein without viral infection (Fig. 6b), and SeV infection strongly stimulated the nuclear translocation of IRF3 in HeLa cells transfected with the control vector (Fig. 6b). However, SeV-induced IRF3 nuclear translocation was significantly decreased when the SARS-CoV-2 M protein was expressed in HeLa cells, compared with the control cells (Fig. 6b, c). Thus, the SARS-CoV-2 M protein exerts its inhibitory function of type I and III IFNs by preventing IRF3 phosphorylation and nuclear translocation.

### The SARS-CoV-2 M protein promotes viral replication

We have shown that the SARS-CoV-2 M protein can suppress type I and III IFN-induced antiviral immunity. Next, it was of interest to determine the role of the SARS-CoV-2 M protein in viral replication. VSV is commonly used as a model virus to study the effect of IFN on viral replication. VSV-eGFP was used to infect HEK293 cells transfected with empty vector or the SARS-CoV-2 M protein plasmid. With fluorescence microscopy, we observed that there were more VSV-eGFP-positive cells in the SARS-CoV-2 M protein-expressing samples than in the control cells transfected with an empty vector (Fig. 7). In the culture supernatant of HEK293 cells expressing the SARS-CoV-2 M protein, the titer of VSV-eGFP was much higher than that of the HEK293 cells transfected with an empty vector (Fig. 7). Moreover, there was more eGFP protein expression in the HEK293 cells expressing the SARS-CoV-2 M protein than in the HEK293 cells transfected with an empty vector (Fig. 7). Thus, the overexpression of the SARS-CoV-2 M protein facilitates the replication of VSV-eGFP.

**Figure 7.**
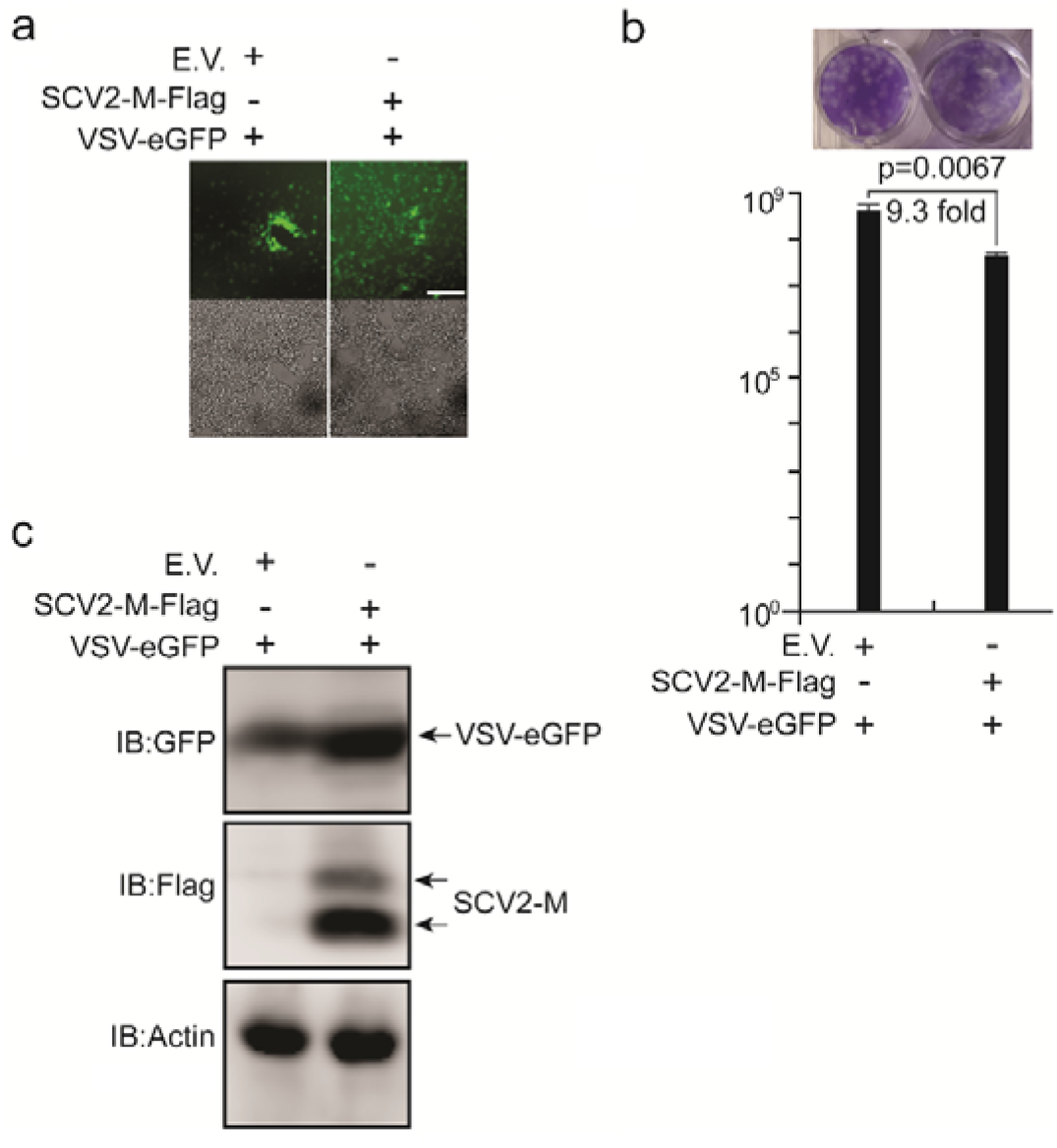
The SARS-CoV-2 M protein facilitates viral replication. The HEK293T cells were transfected with plasmids as indicated, and 24 hours later, the cells were infected with VSV-eGFP (MOI=0.001). Twelve hours after infection, the GFP-positive cells were observed (**a**), and the culture supernatant (20 hours post-infection) was collected for plaque assays to measure the titer of extracellular VSV-eGFP (**b**). Confocal imaging results are representative of two independent experiments. Scale bar, 50 μm. Three independent biological replicates were analyzed; the results of one representative experiment are shown, and the error bars indicate SEM. The statistical significance is shown as indicated. **c** The replication of intracellular VSV-eGFP in the cell lysate (20 hours post-infection) was determined by immunoblotting using an anti-GFP antibody. Empty vector, E.V.; SARS-CoV-2 M protein, SCV2-M.

## Discussion

Our previous studies have shown that innate antiviral immunity may play an important role in the SARS-CoV-2 clearance observed in COVID-19 patients.^38^ The dysregulation of innate antiviral immunity and inflammatory responses by SARS-CoV-2 is strongly responsible for COVID-19-induced human death.^6,39^ Type I and III IFNs are typically suppressed in COVID-19 patients, but the molecular mechanism of this phenomenon caused by SARS-CoV-2 still needs to be elucidated.^6,23^ Here, we reported that the SARS-CoV-2 M protein can target the cytosolic RNA-sensing pathway of RIG-I/MDA-5 signaling to block the activation of type I and III IFN responses and thus inhibit host antiviral immunity.

The M proteins of SARS-CoV-1 and MERS-CoV were reported to suppress type I IFN production by interacting with TRAF3 to disrupt TRAF3-TBK1 association. ^36,37,40^ However, the IFN-antagonizing activities of the MERS-CoV M protein seem to be extremely low. The SARS-CoV-1 M protein interacts with TBK1, but the MERS-CoV M protein cannot be detected to associate with TBK1, suggesting that these two highly pathogenic coronaviruses engage distinctive mechanisms in IFN suppression. Moreover, the M protein of the human coronavirus HKU1 does not impact type I IFN production, indicating that the IFN antagonistic property of the M proteins is not evolutionarily conserved in human coronaviruses.^36,37,40^ Importantly, one recent study found that the SARS-CoV-2 M protein cannot inhibit type I IFN and ISG production.^41^ The differences between the M proteins of these coronaviruses in antagonizing IFN signaling drive us to study the function and mechanism of the SARS-CoV-2 M protein in evading innate antiviral immunity. We first examined the effect of the SARS-CoV-2 M protein on the activation of the cytosolic dsRNA-sensing pathway of RIG-I/MDA-5-MAVS signaling and the endosome dsRNA-sensing pathway of TLR3-TRIF signaling. The results demonstrated that the SARS-CoV-2 M protein can significantly inhibit the activation of type I and III IFN production induced by the RIG-I/MDA-5 signaling components, including RIG-I, MDA-5, MAVS, TBK1, and IKKε, but not by TRIF, the adaptor protein of the TLR3-TRIF signaling, or by STING, the adaptor protein of the cGAS-STING signaling (Fig. 2). These results indicated that the SARS-CoV-2 M protein specifically targets the RIG-I/MDA-5-dependent RNA-sensing pathway but not the TLR3-TRIF-dependent endosome dsRNA-sensing pathway or the cGAS-STING-dependent cytosolic dsDNA-sensing pathway. Furthermore, coimmunoprecipitation studies showed that the SARS-CoV-2 M protein interacts with RIG-I, MDA-5, MAVS, and TBK1 but not with IRF3 (Fig. 4). The overexpression of the SARS-CoV-2 M protein could prevent the complex formation of RIG-I-MAVS, MAVS-TBK1, and TRAF3-TBK1 (Fig. 5). Thus, the SARS-CoV-2 M protein exerts its inhibitory effect on IFN production by impeding the formation of protein complex involving RIG-I, MAVS, TRAF3, and TBK1, which are essential for IRF3 activation and IFN production. We also found that the SARS-CoV-2 M protein can inhibit the phosphorylation of IRF3 but not that of TBK1 (Fig. 6a), which may explain why the SARS-CoV-2 M protein interacts with TBK1 but cannot suppress TRIF- or STING-induced IFN production. This finding also suggests that the SARS-CoV-2 M protein does not directly target TBK1 but acts upstream of TBK1.

Although SARS-CoV-1 and MERS-CoV M proteins were shown to inhibit IRF3 phosphorylation, whether they also affected the translocation of IRF3 into the nucleus was unknown. ^36,37,40^ To extend the understanding of the biological function of the SARS-CoV-2 M protein, we observed that after SeV infection, IRF3 was mainly retained in the cytosol in HeLa cells expressing the SARS-CoV-2 M protein; in addition, in control HeLa cells that did not express the SARS-CoV-2 M protein, IRF3 was dominantly translocated into the nucleus upon SeV infection (Fig. 6b, c). Thus, the SARS-CoV-2 M protein can powerfully inhibit the nuclear localization of IRF3 induced by SeV. Consistently, IRF3 phosphorylation in HeLa cells expressing the SARS-CoV-2 M protein was also attenuated compared with those cells expressing an empty vector (Fig. 6a).

The SARS-CoV-2 M protein possesses three transmembrane motifs (Supplementary Fig. 2). We showed that it could colocalize with markers of the ER and Golgi apparatus (Fig. 3a-c), which are important signaling platforms for innate antiviral immunity.^42,43^ In addition, we observed that the SARS-CoV-2 M protein showed colocalization with TBK1, and partial colocalization with RIG-I, MDA-5, and MAVS but not with IRF3 (Fig. 3d-h; Fig. 6b), and these observations were also consistent with our coimmunoprecipitation results that the SARS-CoV-2 M protein interacted with RIG-I, MAVS, and TBK1 but not with IRF3 (Fig. 4). Moreover, we found that the SARS-CoV-2 M protein impeded the formation of the multi-protein complex of RIG-I, MAVS, and TBK1 (Fig. 5a-d). An interesting question is whether this inhibition results from retention in a cellular compartment that blocks translocalization. A previous study has suggested that the translocalization of TBK1 from the ER or mitochondria to the Golgi fragments is pivotal for SeV-induced IFN activation. ^43^ Since the SARS-CoV-2 M protein is localized in both the ER and the Golgi, its inhibitory effect is likely achieved through inhibiting TBK1 translocation from the ER to the Golgi compartments and subsequently affecting the formation of TBK1-containing Golgi fragments. ^42,43^ Therefore, whether TBK1-containing Golgi fragment formation is affected by the SARS-CoV-2 M protein warrants further analysis, which may represent as a novel mechanism by which virus-encoded proteins render IFN production inefficient.

The M proteins of SARS-CoV-1 and MERS-CoV were reported to inhibit type I IFN production.^36,37,40^ However, the role of the coronavirus M protein in viral replication still needs further investigation. The results from the fluorescence microscopy, plaque assays, and immunoblotting indicated that the replication of VSV-eGFP was significantly enhanced in HEK293T cells expressing the SARS-CoV-2 M protein (Fig. 7). Thus, the SARS-CoV-2 M protein likely promotes viral replication by suppressing the host IFN responses.

Compared with the previous studies on the function of coronavirus M proteins in antagonizing IFNs, we provided several novel findings. First, we showed for the first time that the SARS-CoV-2 M protein suppresses both type I IFN and III IFN production, but the previous studies were restricted to the effect of coronavirus M protein on type I IFN. This finding may explain the advantage of type III IFNs in curing COVID-19.^24^ Second, the SARS-CoV-1 and MERS-CoV M proteins have been reported to impede TRAF3-TBK1 complex formation. In our study, we found that the SARS-CoV-2 M protein not only prevented TRAF3-TBK1 complex formation but also impaired the formation of the multi-protein complex of RIG-I-MAVS-TBK1, which is essential for IRF3 phosphorylation, translocation, and IFN transcription. Third, we also observed that the SARS-CoV-2 M protein specifically inhibited RIG-I/MDA-5-MASV signaling rather than TLR3-TRIF or cGAS-STING signaling, which may provide more precise targets for potential COVID-19 treatment. In addition, we reported that the SARS-CoV-2 M protein inhibited IRF3 phosphorylation and nuclear translocation activated by SeV infection. Last, although the SARS-CoV-1 and MERS-CoV M proteins were reported to suppress type I IFNs, their roles in viral replication are still need to be addressed. In our study, we demonstrated that the overexpression of the SARS-CoV-2 M protein could prevent the replication of VSV-eGFP.

Although we provide ample results to demonstrate that the SARS-CoV-2 M protein inhibits type I and III IFN production, we are aware that the ectopic expression of the viral protein is different from a real infection for the study of its biological function. Since the M protein is a structural protein and indispensable for virion assembly, constructing an M-null SARS-CoV-2 virus is currently not possible. Identification of a mutant of the SARS-CoV-2 M protein that loses IFN suppression activities but does not affect virion assembly merits further investigations. In addition, such a mutant may provide guidelines for the future production of an M-defective mutant virus of SARS-CoV-2, which will help to contribute to our understanding of SARS-CoV-2 M in antagonizing IFN in a real viral infection context.

The administration of type I or III IFNs alone or together with other drugs has resulted in a reduced virus titer, a limited inflammatory response, and mild clinical disease in both animal models and patients infected by SARS-CoV-1 or SARS-CoV-2.^9,24^ Multiple proteins encoded by coronaviruses were shown to counteract the IFN response, although the mechanistic details of their actions for most of these proteins have not been well documented.^8,41^ A better understanding of the viral IFN antagonists in the pathogenesis of SARS-CoV-2 has important implications in the development of new antiviral drugs and vaccines. In this study, for the first time, we studied the function of the SARS-CoV-2 M protein in counteracting the IFN-mediated innate antiviral immunity and its role in viral replication. Our investigation extends the understanding of antiviral immunity evasion strategies employed by SARS-CoV-2 and reveals a novel mechanistic insight on the mechanisms of IFN inhibition by coronavirus M protein, thus shedding light on the interactions between human antiviral immunity and coronavirus infection in the pathogenesis of COVID-19.

## Acknowledgements

This work was supported by grants from the COVID-19 emergency tackling research project of Shandong University (Grant No. 2020XGB03 to P.-H.W), grants from the Natural Science Foundation of Jiangsu Province (SBK2020042706 to P.-H.W), grants from the Natural Science Foundation of China (81930039, 31730026, 81525012) awarded to C.G, and the Fundamental Research Funds of Shandong University (21510078614099), the Fundamental Research Funds of Cheeloo College of Medicine (21510089393109), China Postdoctoral Science Foundation (2018M642662), and the Natural Science Foundation of China (81901604) awarded to Y.Z. We thank Translational Medicine Core Facility of Shandong University for consultation and instrument availability that supported this work

## Contributions

C.G. and P.-H.W. conceptualized the study. Y.Z., M.-W.Z., L.H., and J.Z. performed the experiments. P.-H.W. wrote the first draft of manuscript. All of the authors contributed to revising the manuscript and approved the final version for publication.

---

**Supplemental Table S1.**
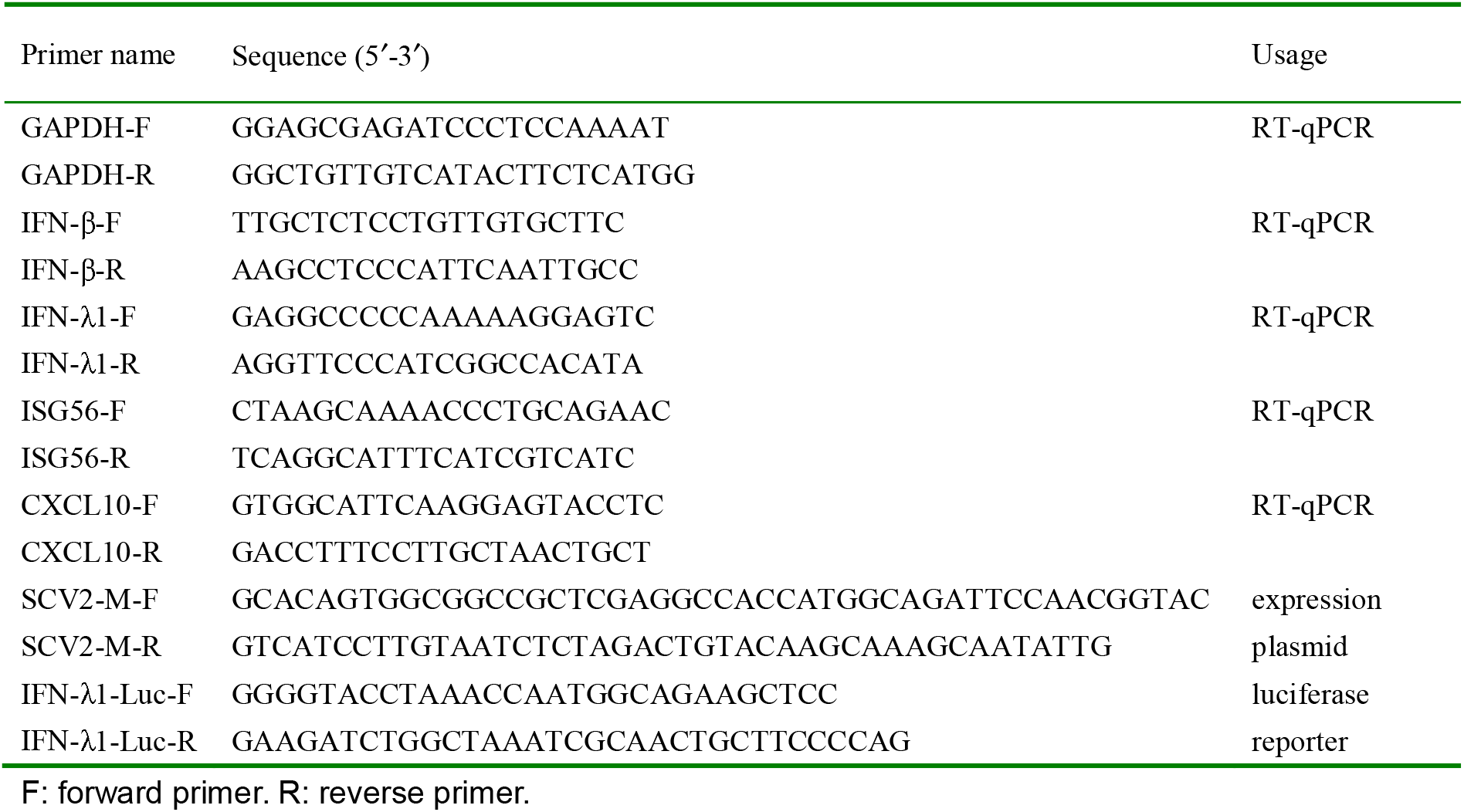
Primers used in this study

**Supplemental Figure 1.**
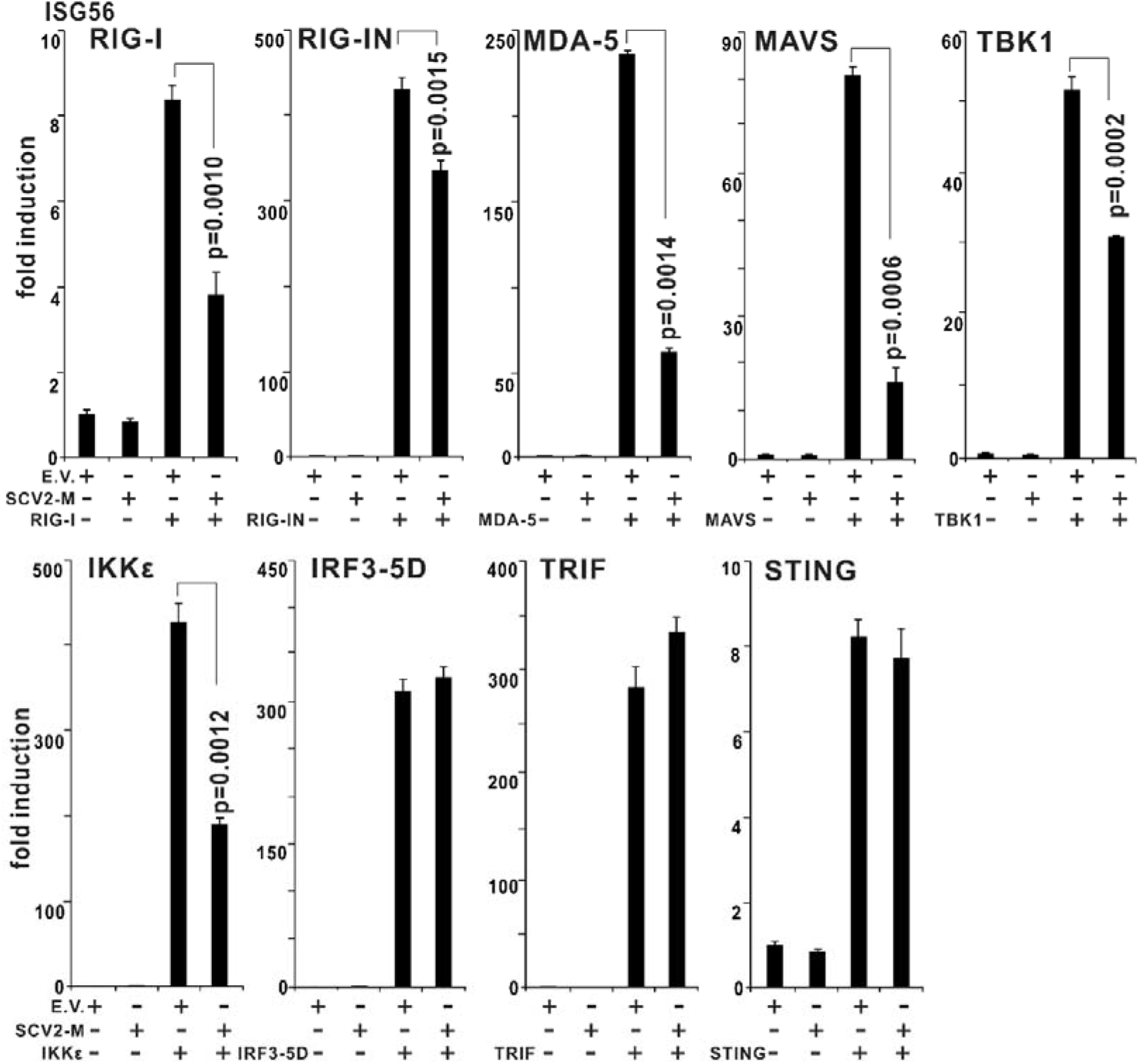
The SARS-CoV-2 M protein impairs IFN-stimulated 56 (ISG56) production induced by the RIG-I/MDA-5 pathway but not by the TLR3-TRIF pathway or the cGAS-STING pathway. The pcDNA6B empty vector (E.V.) and plasmid expressing SARS-CoV-2 M protein (SCV2-M, 300 ng) were transfected with the indicated combinations of plasmids expressing RIG-IN (200 ng), MDA-5 (200 ng), TBK1 (200 ng), IKKε (200 ng), IRF3-5D (200 ng), TRIF (200 ng), or STING (200 ng) into HEK293T cells cultured in 24-well plates (0.8-1 x 10^5^ per well). Thirty-six hours later, the cells were harvested for RNA extraction and subsequent RT-qPCR analysis of the ISG56 induction. Three independent biological replicates were analyzed; the results of one representative experiment are shown, and the error bars indicate SEM. The statistical significance is shown as indicated. Empty vector, E.V.; SARS-CoV-2 M protein, SCV2-M.

**Supplemental Figure 2.**
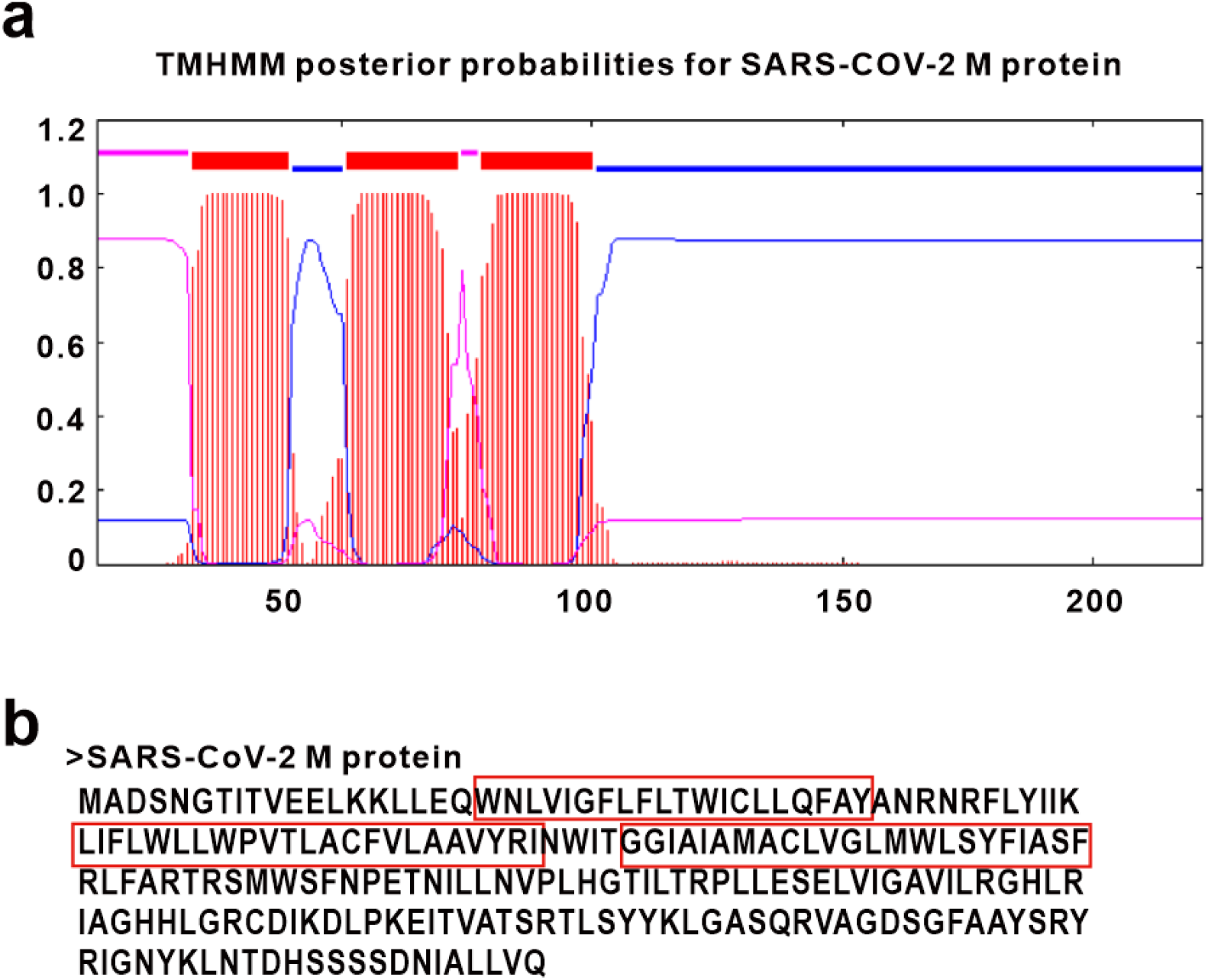
The SARS-CoV-2 M protein is predicated to possess three transmembrane motifs. (A) The transmembrane motifs of the SARS-CoV-2 M protein were predicted with the TMHMM server, version 2.0. (B) The transmembrane motifs of SARS-CoV-2 M protein were labeled with a red box.

## Notes

### Competing Interest Statement

The authors have declared no competing interest.

## References

1 Wu, F. et al. A new coronavirus associated with human respiratory disease in China. Nature 579, 265–269, doi:10.1038/s41586-020-2008-3 (2020).

2 Zhou, P. et al. A pneumonia outbreak associated with a new coronavirus of probable bat origin. Nature 579, 270–273, doi:10.1038/s41586-020-2012-7 (2020).

3 Zhu, N. et al. A Novel Coronavirus from Patients with Pneumonia in China, 2019. N Engl J Med 382, 727–733, doi:10.1056/NEJMoa2001017 (2020).

4 Cui, J., Li, F. & Shi, Z. L. Origin and evolution of pathogenic coronaviruses. Nat Rev Microbiol 17, 181–192, doi:10.1038/s41579-018-0118-9 (2019).

5 Kim, D. et al. The Architecture of SARS-CoV-2 Transcriptome. Cell 181, 914–921 e910, doi:10.1016/j.cell.2020.04.011 (2020).

6 Blanco-Melo, D. et al. Imbalanced Host Response to SARS-CoV-2 Drives Development of COVID-19. Cell 181, 1036–1045 e1039, doi:10.1016/j.cell.2020.04.026 (2020).

7 Ni, L. et al. Detection of SARS-CoV-2-Specific Humoral and Cellular Immunity in COVID-19 Convalescent Individuals. Immunity 52, 971–977 e973, doi:10.1016/j.immuni.2020.04.023 (2020).

8 Totura, A. L. & Baric, R. S. SARS coronavirus pathogenesis: host innate immune responses and viral antagonism of interferon. Curr Opin Virol 2, 264–275, doi:10.1016/j.coviro.2012.04.004 (2012).

9 Park, A. & Iwasaki, A. Type I and Type III Interferons - Induction, Signaling, Evasion, and Application to Combat COVID-19. Cell Host Microbe 27, 870–878, doi:10.1016/j.chom.2020.05.008 (2020).

10 Hu, Y. H. et al. WDFY1 mediates TLR3/4 signaling by recruiting TRIF. EMBO Rep 16, 447–455, doi:10.15252/embr.201439637 (2015).

11 Liu, S. et al. Phosphorylation of innate immune adaptor proteins MAVS, STING, and TRIF induces IRF3 activation. Science 347, aaa2630, doi:10.1126/science.aaa2630 (2015).

12 Sun, L., Wu, J., Du, F., Chen, X. & Chen, Z. J. Cyclic GMP-AMP synthase is a cytosolic DNA sensor that activates the type I interferon pathway. Science 339, 786–791, doi:10.1126/science.1232458 (2013).

13 Onoguchi, K. et al. Viral infections activate types I and III interferon genes through a common mechanism. J Biol Chem 282, 7576–7581, doi:10.1074/jbc.M608618200 (2007).

14 Chang, C. Y., Liu, H. M., Chang, M. F. & Chang, S. C. Middle East Respiratory Syndrome Coronavirus Nucleocapsid Protein Suppresses Type I and Type III Interferon Induction by Targeting RIG-I Signaling. J Virol 94, doi:10.1128/JVI.00099-20 (2020).

15 Yoshikawa, T et al. Dynamic innate immune responses of human bronchial epithelial cells to severe acute respiratory syndrome-associated coronavirus infection. PLoS One 5, e8729, doi:10.1371/journal.pone.0008729 (2010).

16 Hadjadj, J. e. a. Impaired type I interferon activity and exacerbated inflammatory responses in severe Covid-19 patients. medRxiv (2020).

17 Hung, I. F. et al. Triple combination of interferon beta-1b, lopinavir-ritonavir, and ribavirin in the treatment of patients admitted to hospital with COVID-19: an open-label, randomised, phase 2 trial. Lancet 395, 1695–1704, doi:10.1016/S0140-6736(20)31042-4 (2020).

18 Irvani, S. S. N., Golmohammadi, M., Pourhoseingholi, M. A., Shokouhi, S. & Darazam, I. A. Effectiveness of Interferon Beta 1a, compared to Interferon Beta 1b and the usual therapeutic regimen to treat adults with moderate to severe COVID-19: structured summary of a study protocol for a randomized controlled trial. Trials 21, 473, doi:10.1186/s13063-020-04382-3 (2020).

19 Mantlo, E., Bukreyeva, N., Maruyama, J., Paessler, S. & Huang, C. Antiviral activities of type I interferons to SARS-CoV-2 infection. Antiviral Res 179, 104811, doi:10.1016/j.antiviral.2020.104811 (2020).

20 Zhongji Meng, T. W., Chen Li, Xinhe Chen, Longti Li, Xueqin Qin, Hai Li, Jie Luo. An experimental trial of recombinant human interferon alpha nasal drops to prevent coronavirus disease 2019 in medical staff in an epidemic area. medRxiv (2020).

21 Lokugamage, K. G., Hage, A., Schindewolf, C., Rajsbaum, R. & Menachery, V. D. SARS-CoV-2 is sensitive to type I interferon pretreatment. bioRxiv, doi:10.1101/2020.03.07.982264 (2020).

22 Abigail Vanderheiden, P. R., Tatiana Chirkova, Amit A. Upadhyay, Matthew G. Zimmerman, Shamika Bedoya, Hadj Aoued, Gregory M. Tharp, Kathryn L. Pellegrini, View ORCID ProfileAnice C. Lowen, View ORCID ProfileVineet D. Menachery, Larry J. Anderson, Arash Grakoui, Steven E. Bosinger, View ORCID ProfileMehul S. Suthar. Type I and Type III IFN Restrict SARS-CoV-2 Infection of Human Airway Epithelial Cultures. bioRxiv (2020).

23 O’Brien, T R. et al. Weak Induction of Interferon Expression by SARS-CoV-2 Supports Clinical Trials of Interferon Lambda to Treat Early COVID-19. Clin Infect Dis, doi:10.1093/cid/ciaa453 (2020).

24 Stanifer, M. L. et al. Critical Role of Type III Interferon in Controlling SARS-CoV-2 Infection in Human Intestinal Epithelial Cells. Cell Rep, 107863, doi:10.1016/j.celrep.2020.107863 (2020).

25 Davidson, S. et al. IFNlambda is a potent anti-influenza therapeutic without the inflammatory side effects of IFNalpha treatment. EMBO Mol Med 8, 1099–1112, doi:10.15252/emmm.201606413 (2016).

26 Lazear, H. M., Schoggins, J. W. & Diamond, M. S. Shared and Distinct Functions of Type I and Type III Interferons. Immunity 50, 907–923, doi:10.1016/j.immuni.2019.03.025 (2019).

27 Fu Hsin, T.-L. C., Yun-Rui Chan, Han-Chieh Kao, Wang-Da Liu, Jann-Tay Wang, Yu-Hao Pang, Chih-Hui Lin, Ya-Min Tsai, Jing-Yi Lin, Sui-Yuan Chang, View ORCID ProfileHelene Minyi Liu. Distinct Inductions of and Responses to Type I and Type III Interferons Promote Infections in Two SARS-CoV-2 Isolates. bioRxiv (2020).

28 Ziegler, C. G. K. et al. SARS-CoV-2 Receptor ACE2 Is an Interferon-Stimulated Gene in Human Airway Epithelial Cells and Is Detected in Specific Cell Subsets across Tissues. Cell 181, 1016–1035 e1019, doi:10.1016/j.cell.2020.04.035 (2020).

29 Zhuang, M. W. et al. Increasing host cellular receptor-angiotensin-converting enzyme 2 expression by coronavirus may facilitate 2019-nCoV (or SARS-CoV-2) infection. J Med Virol, doi:10.1002/jmv.26139 (2020).

30 Liu, B. et al. The ubiquitin E3 ligase TRIM31 promotes aggregation and activation of the signaling adaptor MAVS through Lys63-linked polyubiquitination. Nat Immunol 18, 214–224, doi:10.1038/ni.3641 (2017).

31 Song, G. et al. E3 ubiquitin ligase RNF128 promotes innate antiviral immunity through K63-linked ubiquitination of TBK1. Nat Immunol 17, 1342–1351, doi:10.1038/ni.3588 (2016).

32 Wang, P. H. et al. A novel transcript isoform of STING that sequesters cGAMP and dominantly inhibits innate nucleic acid sensing. Nucleic Acids Res 46, 4054–4071, doi:10.1093/nar/gky186 (2018).

33 Wang, P. H. et al. Nucleic acid-induced antiviral immunity in shrimp. Antiviral Res 99, 270–280, doi:10.1016/j.antiviral.2013.05.016 (2013).

34 Wang, P. H. et al. The shrimp IKK-NF-kappaB signaling pathway regulates antimicrobial peptide expression and may be subverted by white spot syndrome virus to facilitate viral gene expression. Cell Mol Immunol 10, 423–436, doi:10.1038/cmi.2013.30 (2013).

35 Wang, P. H. et al. Inhibition of AIM2 inflammasome activation by a novel transcript isoform of IF116. EMBO Rep 19, doi:10.15252/embr.201845737 (2018).

36 Siu, K. L. et al. Severe acute respiratory syndrome coronavirus M protein inhibits type I interferon production by impeding the formation of TRAF3.TANK.TBK1/IKKepsilon complex. J Biol Chem 284, 16202–16209, doi:10.1074/jbc.M109.008227 (2009).

37 Lui, P. Y. et al. Middle East respiratory syndrome coronavirus M protein suppresses type I interferon expression through the inhibition of TBK1-dependent phosphorylation of IRF3. Emerg Microbes Infect 5, e39, doi:10.1038/emi.2016.33 (2016).

38 Wang, B. et al. Long-term coexistence of SARS-CoV-2 with antibody response in COVID-19 patients. J Med Virol, doi:10.1002/jmv.25946 (2020).

39 Zhou, Z. et al. Heightened Innate Immune Responses in the Respiratory Tract of COVID-19 Patients. Cell Host Microbe 27, 883–890 e882, doi:10.1016/j.chom.2020.04.017 (2020).

40 Siu, K. L., Chan, C. P., Kok, K. H., Chiu-Yat Woo, P. & Jin, D. Y. Suppression of innate antiviral response by severe acute respiratory syndrome coronavirus M protein is mediated through the first transmembrane domain. Cell Mol Immunol 11, 141–149, doi:10.1038/cmi.2013.61 (2014).

41 Yuen, C. K. et al. SARS-CoV-2 nsp13, nsp14, nsp15 and orf6 function as potent interferon antagonists. Emerg Microbes Infect 9, 1418–1428, doi:10.1080/22221751.2020.1780953 (2020).

42 Tao, Y., Yang, Y., Zhou, R. & Gong, T. Golgi Apparatus: An Emerging Platform for Innate Immunity. Trends Cell Biol 30, 467–477, doi:10.1016/j.tcb.2020.02.008 (2020).

43 Pourcelot, M. et al. The Golgi apparatus acts as a platform for TBK1 activation after viral RNA sensing. BMC Biol 14, 69, doi:10.1186/s12915-016-0292-z (2016).

